# The Impact Of Thigh And Shank Marker Quantity On Lower Extremity Kinematics Using A Constrained Model

**DOI:** 10.1101/290890

**Authors:** Annelise A Slater, Todd J. Hullfish, Josh R. Baxter

**Author notes:** Corresponding author: Josh R. Baxter, PhD;, Mailing address: 3737 Market Street, Suite 702, Philadelphia, PA, USA 19104.

## Abstract

Musculoskeletal models are commonly used to quantify joint motions and loads during human motion. Constraining joint kinematics simplifies these models but the implications of the number of markers used during data acquisition remains unclear. The purpose of this study was to establish the effects of marker placement and quantity on kinematic fidelity when using a constrained-kinematic model. We hypothesized that a constrained-kinematic model would faithfully reproduce lower extremity kinematics regardless of the number of tracking markers removed from the thigh and shank. Healthy-young adults (N = 10) walked on a treadmill at slow, moderate, and fast speeds while skin-mounted markers were tracked using motion capture. Lower extremity kinematics were calculated for 256 combinations of leg and shank markers to establish the implications of marker placement and quantity on joint kinematics. Sagittal joint and hip coronal kinematics errors were smaller than documented errors caused by soft-tissue artifact, which tends to be approximately 5 degrees, when excluding thigh and shank markers. Joint angle and center kinematic errors negatively correlated with the number of markers included in the analyses (R^2^ > 0.97) and typically showed the greatest error reductions when two markers were included. Further, we demonstrated that a simplified marker set that included markers on the pelvis, lateral knee condyle, lateral malleolus, and shoes produced kinematics that strongly agreed with the traditional marker set. In conclusion, constrained-kinematic models are resilient to marker placement and quantity, which has implications on study design and post-processing workflows.

***Ethics Approval and Consent to Participate*** this study was approved by the Institutional Review Board at the University of Pennsylvania (#824466). Subjects provided written-informed consent

***Consent to Publish*** this submission does not contain any individual data

***Availability of Data and Materials*** the datasets analyzed in this study are available from the corresponding author on reasonable request.

***Competing Interests*** one author (JB) is an associate editor for BMC Musculoskeletal Disorders. None of the other authors have any competing interests.

***Funding*** no funding has been provided for this research

**Authors’ Contributions:** AS, TH, and JB designed the experiment; AS and TH collected the data; AS and JB analyzed and interpreted the data; AS and JB drafted the manuscript; AS, TH, and JB revised the intellectual content of the manuscript; AS, TH, and JB approved the final version of the manuscript; and AS, TH, and JB agreed to be accountable for all aspects of the study.

## Introduction

Musculoskeletal modeling relies on accurate experimental data to calculate the motions and loads generated during human motion. Despite recent advances in motion capture technology that have improved marker tracking to sub-millimeter precision, soft-tissue artifact continues to be a major limiter of the clinical efficacy of motion capture data [1]. A recent special edition of the Journal of Biomechanics proposed new and innovative techniques to mitigate some of the effects of soft-tissue artifact [2]. While these techniques improve the overall fidelity of motion capture data, they introduce new challenges to both the collection and processing workflows [3–7]. This study takes a different approach to the problem; instead, seeking to understand how currently implemented techniques can be streamlined to preserve tolerable fidelity compared to unconstrained-kinematic models while reducing the burdens placed on subjects and researchers.

Unconstrained-kinematic models – often referred to as ‘six degree-of-freedom’ – are commonly utilized to quantify joint motion using skin-based motion capture [8,9]; however, their accuracy has been challenged by recent fluoroscopy and bone-pin studies [10,11]. For example, knee valgus and internal rotation errors of 117 and 192%, respectively, have been reported despite utilizing techniques that are aimed at minimizing soft tissue artifact [12]. In addition, unconstrained joints increase the complexities of musculoskeletal models, making simulation of human motion challenging.

Constrained-kinematic models leverage well-known characteristics of joint function [13,14] to compensate for soft-tissue artifact while minimizing the number of markers needed to quantify motion [15]. These models also make possible advanced analyses of neuromuscular function and forward dynamic simulations [16] without the need of simulating joint contact, which is impractical to implement on large data sets. Despite these inherent strengths of constrained-kinematic models, experimental considerations of marker placement and quantity have not yet been associated with kinematic fidelity.

The purpose of this study was to rigorously characterize the implications of marker placement and quantity on kinematic fidelity using a constrained-kinematic model. To do this, we tested 256 combinations of marker number and placement and characterized their effects on lower extremity kinematics and joint centers – a surrogate measure of joint kinetics [17]. We hypothesized that (1) joint kinematics calculated using constrained and unconstrained paradigms would not differ and (2) lower extremity kinematic errors would positively correlate with the number of markers excluded from the analyses. The secondary aim of this study was to identify a ‘simplified’ marker set that provides kinematic fidelity while minimizing the number of markers needed for model definition and kinematic tracking.

## Methods

Motion analysis was performed on 10 healthy-young adults (24 ± 4 years, 6 females, BMI 24.2 ± 3.4) who provided written consent in this IRB approved study. Retro-reflective markers (9.5 mm, B&L Engineering, Santa Ana, CA) were placed on the lower-extremities of each subject and tracked using a 12-camera motion capture system (Raptor Series, Motion Analysis Corp, Santa Rosa, CA) while subjects walked on a treadmill (TMX428, Trackmaster, Newton, KS). Markers were placed over anatomic landmarks (Figure 1) of the pelvis: anterior and posterior superior iliac spines; legs: lateral knee condyle and lateral ankle malleolus; and feet: calcaneus, first and fifth metatarsal heads, and the great toe that were placed on the shoes. Additional tracking markers were placed on the proximal-lateral, distal-lateral, and middle-anterior regions of the thigh and shank [18]. Marker positions were acquired while subjects stood in a neutrally-aligned position, which were used to scale a generic musculoskeletal model. Next, subjects walked on a treadmill at a slow (0.9 m/s), moderate (1.2 m/s), and fast (1.5 m/s) pace. Each trial lasted 2 minutes and generated approximately 100 strides for each leg.

**Figure 1.**
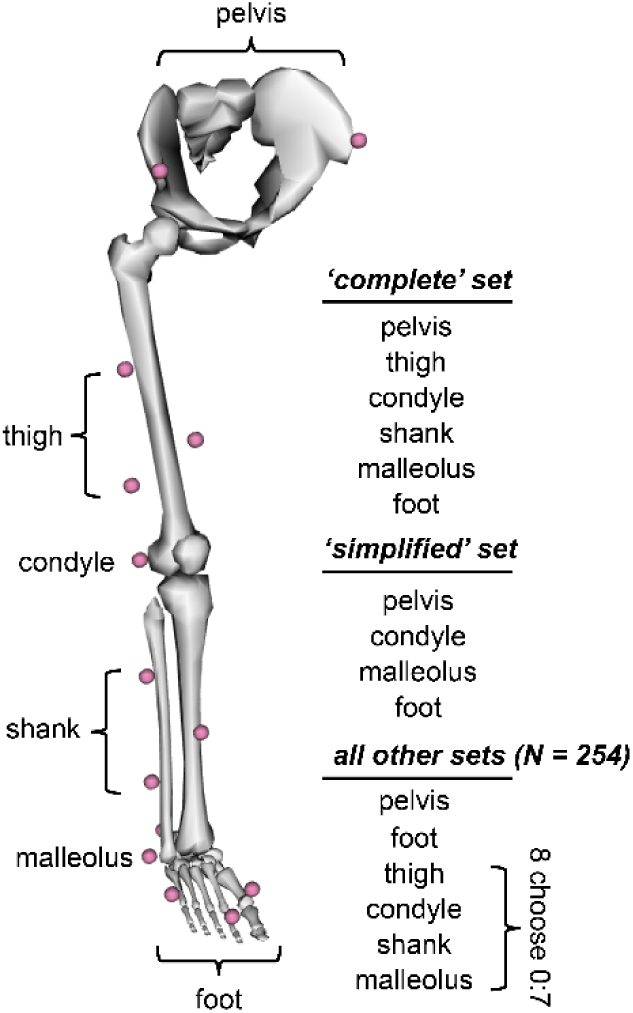
Subject-specific models (left leg hidden for clarity) were scaled based on subject anatomy and positioning. Inverse kinematics were then performed for walking trials under 256 marker combinations to test the effects of all possible marker positions and quantities attached on the thigh and shank.

### Constrained-kinematic model

Lower extremity kinematics were calculated for 256 different combinations of thigh asd shank markers using a constrained-kinematic model implemented in open-source musculoskeletal modeling software (Opensim v3.3; [19]). This lower extremity model [18] – defined the hip as a ball joint, the knee as a mobile-hinge joint, the foot and ankle as an oblique universal joint, and the forefoot as a hinge joint – was scaled based on anatomic landmarks captured in the neutrally-aligned position (Supplemental Material). We used this single degree-of-freedom knee joint that prescribed non-sagittal motions [13] for two reasons: 1 – soft-tissue artifacts cause errors greater in magnitude than the actual joint motion in the coronal and transverse planes [10,20,21] and 2 – the muscles that cross the knee joint do have limited leverage outside of the sagittal plane. Marker trajectories were interpolated using a cubic-spline and low-pass filtered at 6 Hz [8]. Hip, knee, and ankle kinematics were calculated using an inverse kinematics paradigm and all markers received equal weighting [19]. Markers on the thigh and shank segments were systematically excluded from the analysis (Figure 1), resulting in 256 marker combinations tested to characterize the effects of marker location and inclusion on joint kinematics.

Subject-specific musculoskeletal models were scaled using a previously reported generic model [18] and marker positions captured while subjects stood in the anatomic position. The pelvis, thighs, shanks, and feet were scaled based on markers placed on anatomic landmarks: pelvis – right and left anterior superior iliac spines; thigh – anterior superior iliac spine and lateral condyle; shank – lateral condyle and lateral malleolus; and foot – lateral malleolus and toe. The scaled model was then moved to the anatomic position by fitting the model to the anatomic marker positions and recorded joint angles. The anterior superior iliac spines, lateral condyles and malleoli, heel, 1^st^ and 5^th^ metatarsal heads, and toe markers were all given equal weighting. Similarly, the hips, knees, ankles, and toe joints were all weighted towards neutral sagittal alignments. Since hip adduction and rotation varied amongst subjects during the anatomic pose, those coordinates received no weighting. Finally, scaled models were confirmed by superimposing the marker positions over the model.

During the pilot testing for this study (N = 3), we calculated the functional hip joint centers [22] and compared these locations to the hip joint centers from the scaled models [18]. We found that the functional hip joint centers were 30% wider than the generic model, which agrees with prior reports of pelvic morphology [23]. Therefore, we increased the hip joint center width in the unscaled generic model and scaled this modified model for all research subjects based on pelvis anatomy. This had appreciable effects on the initialization of models during pilot testing, where the model positioning agreed more strongly with the marker positions when the wider hip joint center locations were implemented. In order to show the robustness of the constrained-kinematic model, we chose not to modify the hip joint center locations based on subject-specific functional hip joint locations. However, hip kinetics are sensitive to joint center location and employing more rigorous scaling techniques should be considered when high-fidelity hip kinetics are required.

### Unconstrained-kinematic model

Unconstrained joint kinematics were calculated to confirm if the unconstrained and constrained calculations yielded similar results. Anatomic coordinate systems were assigned to each segment using established definitions [24,25] that mirrored the coordinate systems defined in the constrained-kinematic model. Four markers on each segment were used to track and define joint motions and a least squares approach was implemented to minimize the effects of soft-tissue artifact [26]. Euler rotations were calculated in a flexion-adduction-rotation sequence, and joint angles during the anatomic pose trial were matched with the joint angles calculated in the constrained-kinematic model in order to perform a true one-to-one comparison.

### Joint kinematics analysis

Joint angles and centers during each of the 100 measured strides were averaged over each walking speed and marker combination then compared to the kinematics calculated from the complete marker set. Heel strike events were identified using a kinematic-based algorithm [27]. Maximal and minimal joint rotations as well as joint range of motion were calculated for hip flexion and adduction as well as knee flexion and ankle plantarflexion. Cross-correlation coefficients [28] and root mean square (RMS) errors were calculated for joint kinematics. Ninety-five percent confidence intervals were calculated using a bootstrap approach [29] to demonstrate the amount of certainty in the joint kinematics and visualized in plots. Two primary analyses were performed: 1 – joint angles calculated using the unconstrained and constrained models that included all tracking markers and 2 – joint angles and centers for each marker combination using the constrained-kinematic model were compared to the constrained model that included all tracking markers. Joint center displacements in the anterior-posterior, superior-inferior, and medial-lateral directions were calculated with respect to joint center positions from the complete marker set. Prior to data analysis, we defined a ‘substantially different’ cross correlation coefficient (*r*_*xy*_) to be less than 0.9. Hip internal rotations were also calculated as part of a secondary analysis.

Joint kinematics calculated at three walking speeds were compared to determine if a ‘simplified’ marker set – consisting of markers on the pelvis, lateral condyles, lateral malleoli, and shoes – detects speed-dependent changes in joint excursions similarly to a traditional marker set. This simplified marker set was selected because it is easily implemented and the markers placed on the lateral knee and ankle joints are needed to initialize the musculoskeletal model. Group means were compared using paired t-tests and corrections of multiple comparisons were not performed to decrease the likelihood of type II errors, thus making these analyses less conservative and more likely to reject the null hypothesis (no difference between marker sets) when a difference exists.

### Accounting for uncertainty associated with soft-tissue artifact

Lower extremity kinematics derived from skin-mounted markers vary approximately 5 degrees from bony motion [1,20]. Thus, differences in peak joint rotations and range of motions less than this 5 degrees threshold were considered to be within the uncertainty threshold and were not considered for statistical testing [20,30]. Paired t-tests were performed on instances in which this 5 degree thresholds were exceeded to test for statistically significant differences (*p* < 0.05). These bootstrapped confidence intervals calculated from the complete marker set data were expanded by 5 degrees to demonstrate the uncertainty associated with skin mounted markers compared to more direct techniques [1,20].

## Results

Constrained and unconstrained-kinematic paradigms showed strong agreement for sagittal joint motions but moderate agreement for hip adduction (RMS errors: 1.6 – 3.2°; Figure 2). Hip and knee flexion patterns were strongly correlated (*r*_xy_ ≥ 0.90), ankle sagittal motions fell just below the cutoff value for ‘substantially different’ (0.85 < *r*_xy_ < 0.90). Hip adduction patterns were moderately correlated (0.65 < *r*_xy_ < 0.71). Despite any detected differences in kinematic patterns, joint excursions varied by less than five degrees between unconstrained and constrained paradigms.

**Figure 2.**
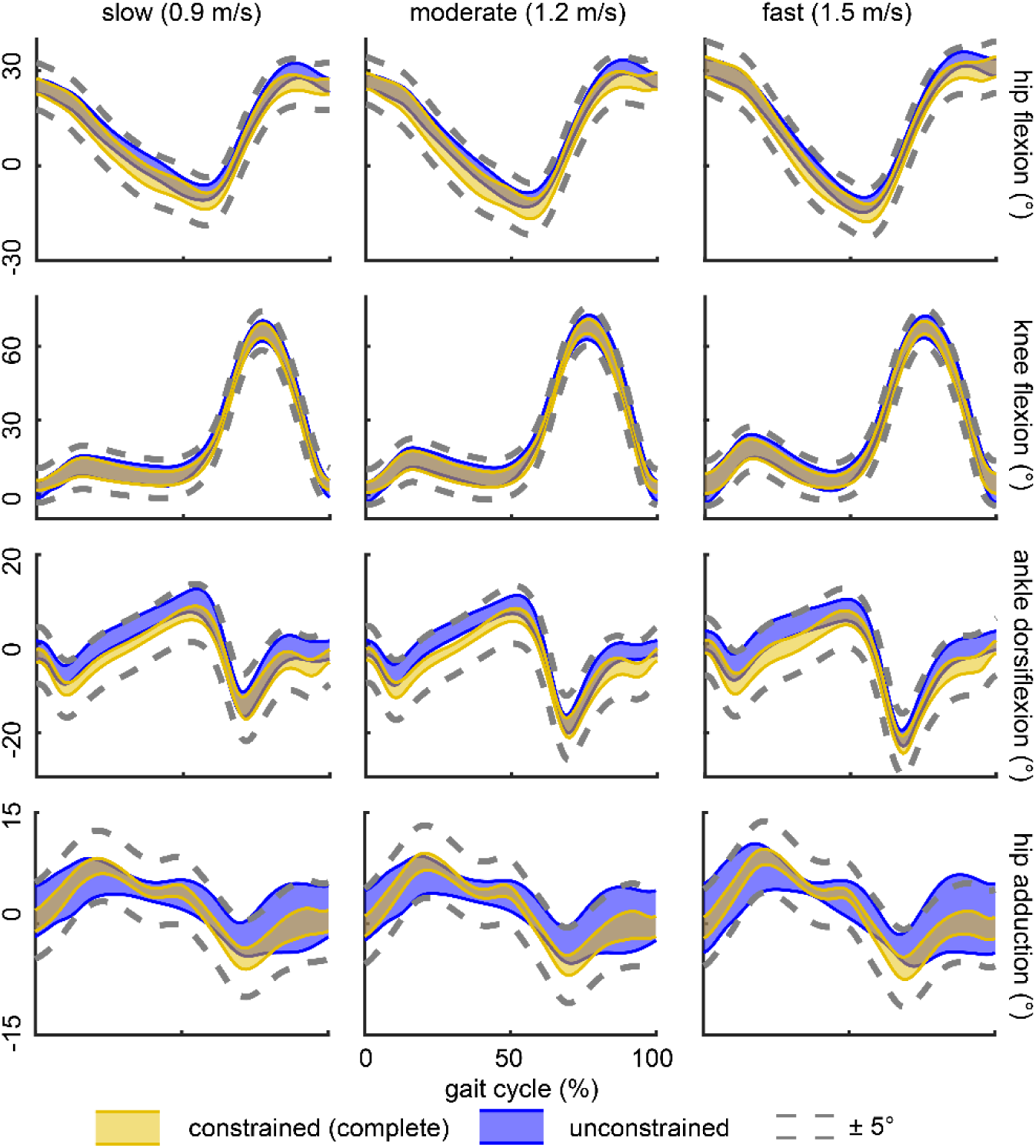
Joint kinematics strongly agreed (*r*_xy_ ≥ 0.90) between the complete marker set (gold band) and all 255 other marker combinations (gray band). The 5° uncertainty threshold (dashed lines) was not exceeded by any marker combination. Specifically, the ‘simplified’ marker set (red band) produced similar joint motions at all three walking speeds. 95% confidence intervals were calculated using a bootstrapping technique to characterize the variability within our study cohort (N = 10).

Lower extremity sagittal kinematics were not strongly affected by removing thigh and shank markers from the kinematic analysis (Figure 3). Specifically, including markers on the lateral knee condyles and malleoli generated high-fidelity sagittal kinematics compared to the constrained-kinematic model that utilized all tracking markers (*r*_xy_ ≥ 0.94; RMS errors < 2.3°). Regardless of the number of markers included in the kinematic analyses, adduction patterns were similar (0.85 < *r*_xy_ < 0.90) and joint range of motion as well as flexion and extension peaks did not deviate beyond the 5° uncertainty threshold. Hip adduction was accurately measured by all but two marker sets – when all markers proximal of the lateral malleoli were removed.

**Figure 3.**
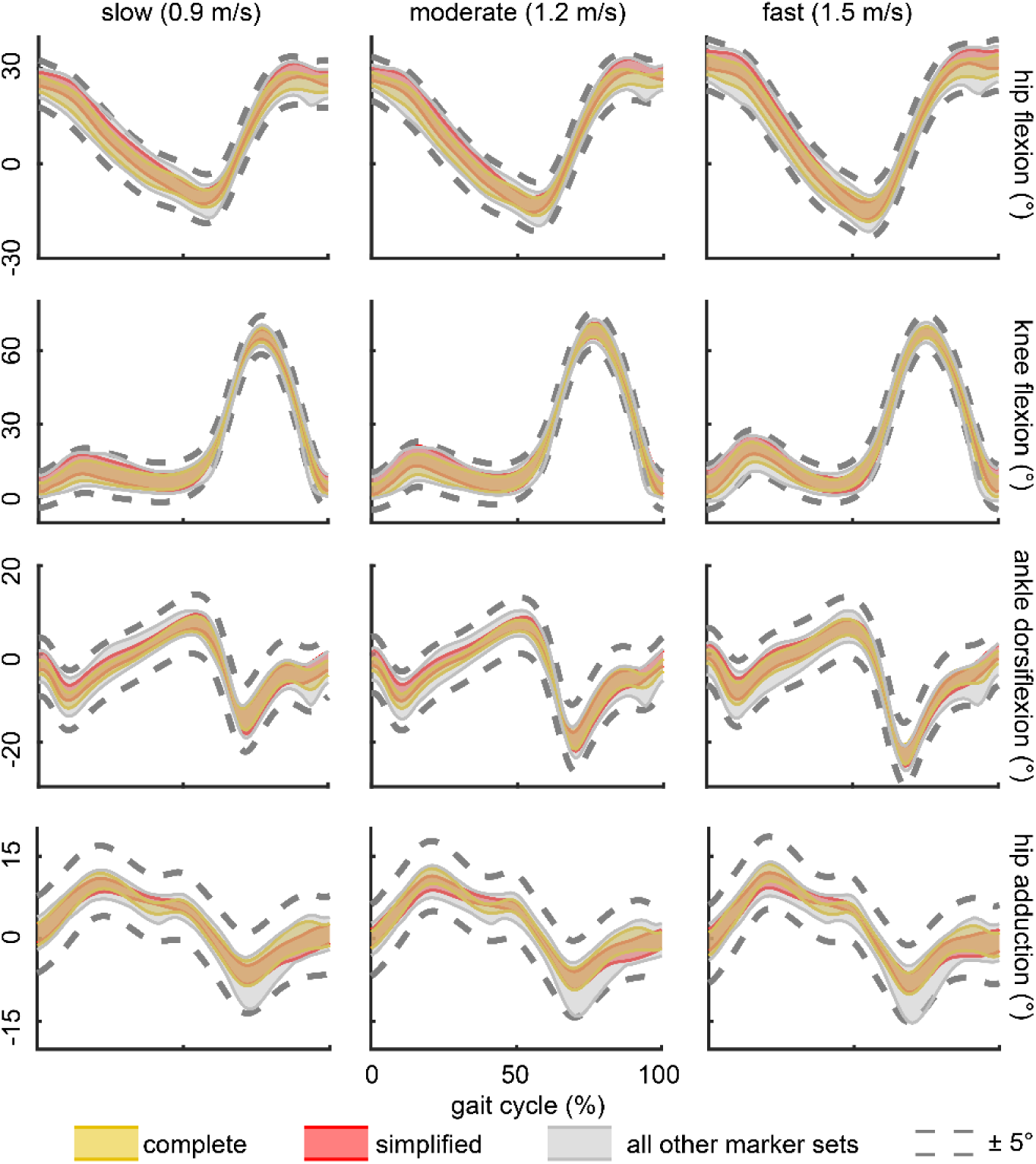
Lower extremity kinematics strongly agreed (*r*_xy_ ≥ 0.90) between the complete marker set (gold band) and all 255 other marker combinations (gray band). The 5° uncertainty threshold (dashed lines) was not exceeded by any marker combination – including the ‘simplified’ marker set (red band) – in the sagittal plane and only 2 marker sets resulted in differences in hip adduction excursion greater than 5°. 95% confidence intervals were calculated using a bootstrapping technique to characterize the variability within our study cohort (N = 10).

Joint angle and center kinematic errors were negatively correlated with the number of markers included in the constrained-kinematic analysis (Figure 4). Joint angle errors decayed at a rate that was strongly non-linear (R^2^ > 0.97, Figure 4), where most of the errors were reduced by including two markers placed on either the thigh or shank in the kinematic analyses. Knee joint center errors were 2-4 fold greater than hip and ankle joint center errors, respectively. Including additional markers in the kinematic analyses had a strong-linear effect (R^2^ > 0.97, Figure 4) on hip and ankle joint center errors, while knee joint center errors decayed at a cubic rate (R^2^ = 0.99, Fig 3B) with diminishing improvements after two markers were included. Hip and ankle joint center positions were less affected by reduced markers (RMS error < 6 mm) than the knee joint (RMS error < 19 mm).

**Figure 4.**
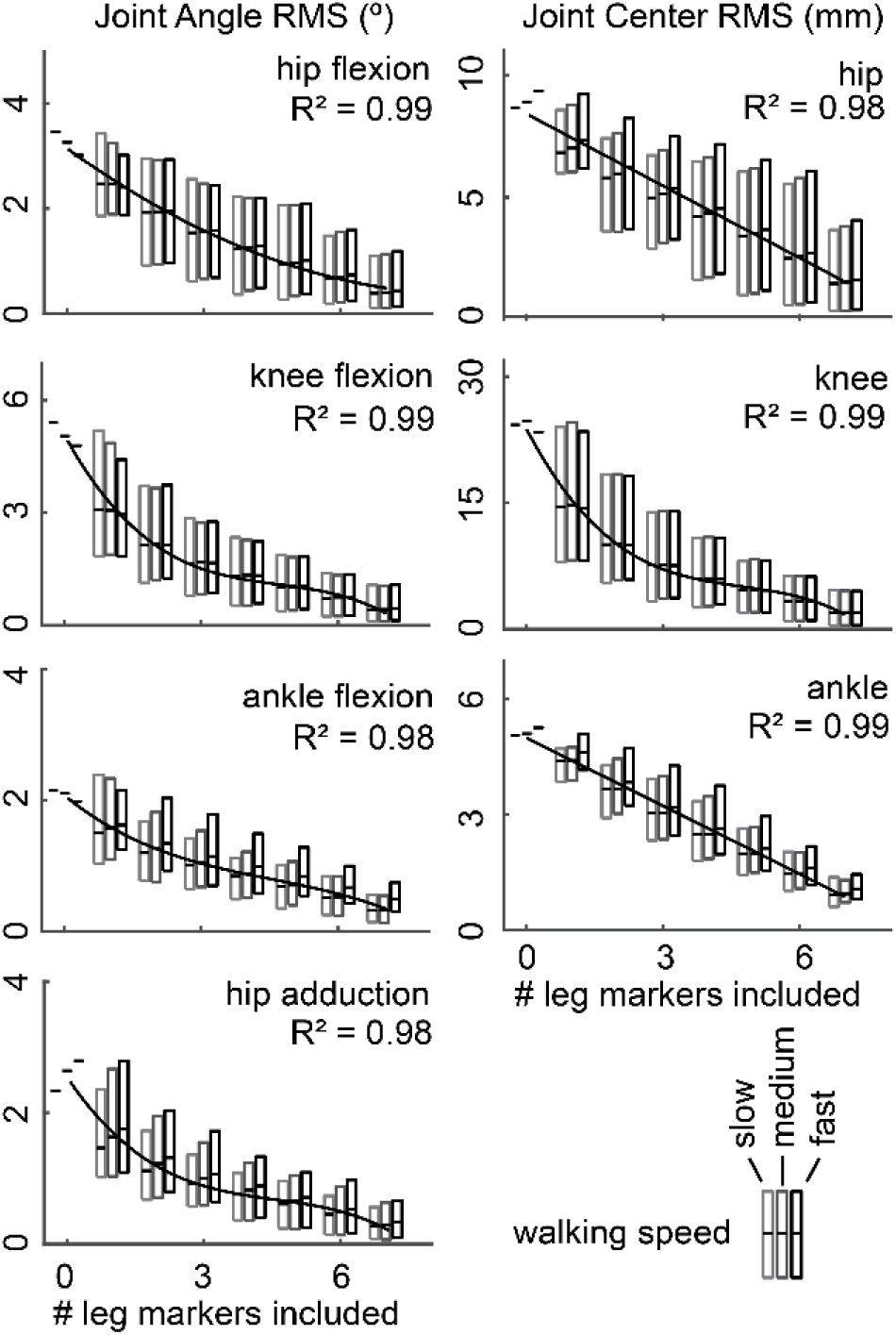
Lower extremity RMS errors negatively correlated with the number of leg markers included in the kinematic analyses. Joint kinematics (angles; left column) errors were best described by a 3^rd^ order polynomial (R^2^ > 0.98). Knee center errors were also best characterized by a 3^rd^ polynomial (R^2^ = 0.99; right column), while hip and ankle center errors linearly correlated with the number of leg markers included in the analyses (R^2^ > 0.98; right column). Walking speed did not affect kinematic errors.

Stereotypical increases in joint excursions were identified with both the complete and simplified marker sets (Table 1; Figure 2). The complete and simplified marker sets demonstrated similar fidelity in detected increases in sagittal joint excursion. Similarly, hip adduction increased with walking speed; however, subtle increases of less than 2° between moderate and fast walking speeds were only detected with the complete marker set.

**Table 1.**
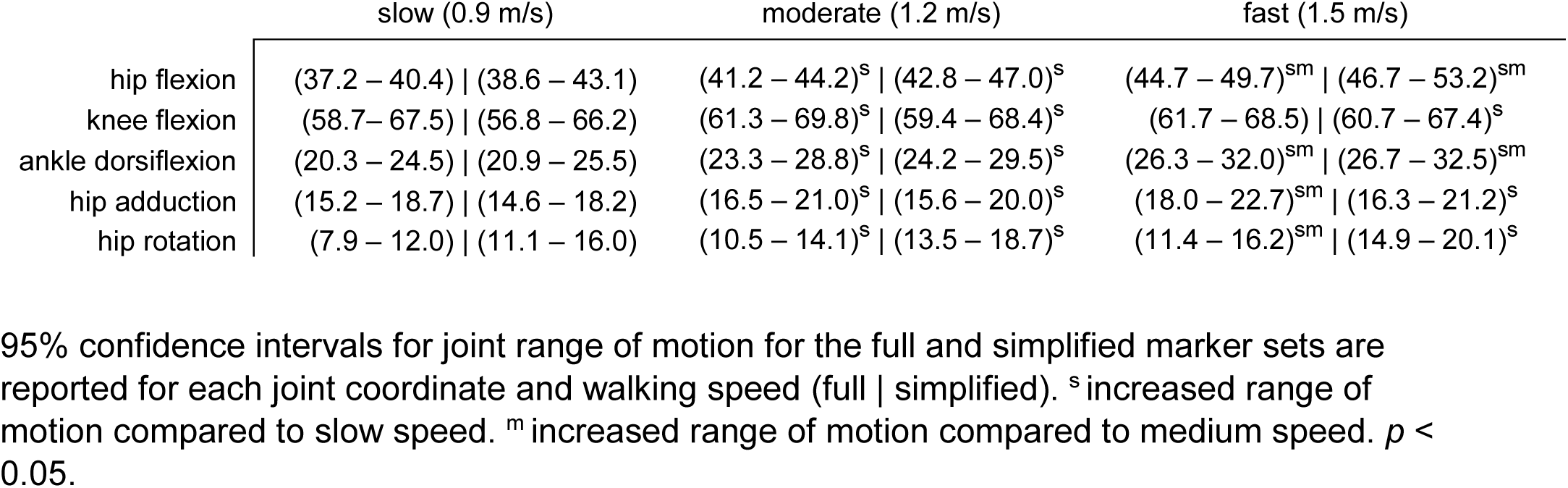
95% confidence intervals for lower extremity ranges of motion – calculated using the full and simplified marker sets – during walking at increasing speeds.

Hip internal rotation patterns were weakly correlated (0.10 < r_xy_ < 0.21) with calculations using an unconstrained-kinematic model and demonstrated differences that exceed five degrees (RMS errors: 3.9 – 5.4°). The effects of removing thigh and shank markers from the constrained-kinematic model had moderate effects (0.55 < *r*_xy_ < 0.90). However, hip internal rotation excursions were within five degrees of the complete marker set in 95% of the marker combinations.

## Discussion

We demonstrated that constrained-kinematic models accurately reproduce lower extremity kinematics when numerous markers are excluded from the analyses. The effects of reducing markers on sagittal kinematics and hip adduction are smaller than kinematic uncertainty caused by soft tissue artifact [1,20,21]. Joint center trajectories, which govern the joint moment arm of the ground reaction force – and thus joint kinetics (Myers, 2015) – appear to also be resilient to decreased markers. Since marker placement minimally affects joint kinematics, researchers can tailor marker sets based on experimental constraints. For example, a ‘simplified’ marker set (Figure 1), that excludes the traditional tracking markers adhered to the thigh and shank, can be utilized without compromising kinematic fidelity to increase motion capture workflow and provide more flexibility for the placement of other sensors and wearable devices.

Lower extremity kinematics quantified in this study compared favorably with prior reports. Similar to prior work [31,32], we found that sagittal hip, knee, and ankle excursion increased with walking speed (Table 1). Hip coronal kinematics measured in this study demonstrated stereotypical patterns that are well described in the literature [33,34]. Since much of the literature implements six degree-of-freedom marker sets, we calculated the unconstrained motion of the lower extremity and implemented a least squares approach [26] to minimize the effects of soft tissue artifact on resulting joint kinematics. Sagittal joint and hip coronal motions were similar between the unconstrained and constrained-kinematic results (Figure 2). Hip internal rotation differed between the unconstrained and constrained model, which may be explained by well documented soft tissue artifact of the thigh segment [35]. However, these differences were less pronounced between constrained marker sets, likely due to the lack of knee rotation in the musculoskeletal model.

Our findings demonstrate that constrained-kinematic models are resilient to marker placement and dropout. Although we did not directly track skeletal motion in this study, we approximated the uncertainty introduced by soft-tissue artifact by calculating the difference between experimentally-measured marker and model-fixed marker trajectories. We found that the markers placed on the thigh, lateral knee, and shank had average RMS values of 12.5, 14.1, and 7.1 mm, respectively; compared to direct measurements of soft-tissue artifact in the literature of 13.8, 13.9, and 10.8 mm, respectively [20]. Despite the lateral knee being prone to soft-tissue artifact, its inclusion improved kinematic tracking when fewer than five leg markers were included in the kinematic analyses (Figure 5). Increasing the number of markers used for gait analysis has diminishing returns with regard to lower extremity kinematics (Figure 4).

**Figure 5.**
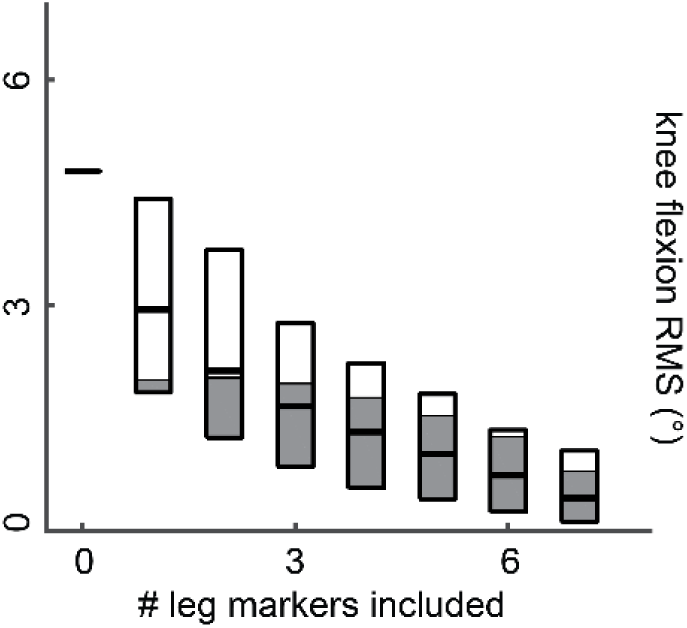
Including the lateral knee condyle marker (shaded part of box) in the constrained-kinematic model effectively decreased the kinematic errors compared to marker sets that excluded the knee marker (unshaded part of box).

Excluding all of the markers attached to the thigh and shank generated sagittal joint kinematics that were in strong agreement with the complete marker set but adversely affected knee joint center kinematics, which impacts joint loads [17]. Adding markers to the lateral condyles and malleoli – which were used to scale the musculoskeletal model – mitigated the majority of kinematic errors (*r*_xy_ ≥ 0.94; RMS errors < 2.3°). To improve experimental consistency and workflow, markers can be permanently fixed to lab shoes, which reduces the number of markers applied to the subject to eight: four on the pelvis and two on each leg. Thus, a ‘simplified’ marker set accurately characterizes joint kinematics and joint center motions by providing essential inputs to constrained-kinematic models.

Minimizing the tracking markers reduces experimental setup time, prevents errors from poorly placed and inherently noisy markers, provides fewer obstructions for other experimental equipment, and allows for more comfortable attire to be worn during data collection. While unconstrained-kinematic models require at least three markers on each segment at all times, our results demonstrate that constrained-kinematic models can perform well with no markers on certain segments; for example, the thigh and shank. Hierarchical marker sets track segment kinematics by assuming the location of a joint center based on a nearby segment. However, this approach is susceptible to soft-tissue artifact [36] and does not provide the necessary joint constraints for advanced musculoskeletal analyses [37]. Our findings also benefit researchers utilizing wearable-assistive devices [38,39], ultrasonography during human motion [40], and high-density electromyography sensors [41] – all techniques that require unobstructed access to the lower extremities.

Processing and analyzing motion capture data can be streamlined into a turn-key routine utilizing open-source musculoskeletal modeling software [19] and batched scripts. In addition to calculating joint kinematics, constrained-kinematic models are well-suited for performing both inverse and forward dynamic simulations. Integrating gait analysis into a single musculoskeletal modeling environment provides investigators with a standardized workflow while maintaining the flexibility needed to perform specific analyses [42,43]. Further, many analyses are not possible to perform without imposing joint constraints or contact [44,45]. Therefore, migrating kinematic analyses into a constrained-kinematic paradigm may minimize workflow complexity without compromising kinematic fidelity (Figure 2).

Several limitations should be considered when interpreting these findings. We did not directly measure skeletal motion but did show similarities in joint kinematics with prior studies that utilized intracortical bone pins and fluoroscopy [20,46]. Subjects in the present study were healthy-young adults that were generally fit with a healthy body mass index (BMI 24.2 ± 3.4), which may not be representative of clinical populations. While it is expected that soft-tissue artifact increases with excess tissue [47] and high-impact activities, constrained-kinematic models appear to be well suited to minimize such effects. Joint kinematics are sensitive to joint-axis location and orientation [48,49], which may be affected when scaling generic musculoskeletal models to subject-specific anthropometry. To mitigate these potential errors, we visually confirmed that each subject-specific model closely matched the neutrally-aligned position. Further, we confirmed joint kinematics using unconstrained-kinematic models that shared the same joint axis definitions as the constrained-kinematic models (Figure 2). Due to knee valgus and internal rotation errors as high as twice that of skeletal motion [12], we limited knee joint kinematics to a single degree-of-freedom and prescribed other rotations and translations based on flexion angle [13]. Walking trials were acquired on a commercial treadmill that did not have an integrated force plate, so we were unable to calculate joint reaction moments. We instead decided to quantify the changes in the joint center trajectories, which governs the ground reaction force moment arm and thus joint moments.

In conclusion, constrained-kinematic models are resilient to errors caused by marker placement and quantity. Experiments can be designed to attain the kinematic fidelity necessary to answer specific research questions while adjusting marker placement and quantity to suit the constraints of the experimental setup. In addition, integrating constrained-kinematic models into a gait analysis workflow offers several advantages that can improve post-processing efficiency while providing access to unique analysis tools to test specific questions. However, investigators should weigh the strengths and weaknesses of both constrained and unconstrained-kinematic paradigms to determine which approach is best suited for the specific research question.

## Acknowledgements

the Authors have no acknowledgements

## References

1. Leardini A, Chiari L, Della Croce U, Cappozzo A. Human movement analysis using stereophotogrammetry. Part 3. Soft tissue artifact assessment and compensation. Gait Posture. 2005 Mar;21(2):212–25.

2. Camomilla V, Dumas R, Cappozzo A. Human movement analysis: The soft tissue artefact issue. J Biomech. 2017 Sep 6;62:1–4.

3. Barré A. Assessment of the lower limb soft tissue artefact at marker-cluster level with a high-density marker set during walking. 2017;

4. Begon M, Bélaise C, Naaim A, Lundberg A, Chèze L. Multibody kinematics optimization with marker projection improves the accuracy of the humerus rotational kinematics. J Biomech. 2016 Oct;

5. Masum MA, Pickering MR, Lambert AJ, Scarvell JM, Smith PN. Multi-slice ultrasound image calibration of an intelligent skin-marker for soft tissue artefact compensation. J Biomech. 2017 Sep 6;62:165–71.

6. Richard V, Cappozzo A, Dumas R. Comparative assessment of knee joint models used in multi-body kinematics optimisation for soft tissue artefact compensation. J Biomech. 2017 Sep 6;62:95–101.

7. Sangeux M, Barré A, Aminian K. Evaluation of knee functional calibration with and without the effect of soft tissue artefact. J Biomech [Internet]. 2016 Nov [cited 2017 Sep 12]; Available from: http://linkinghub.elsevier.com/retrieve/pii/S0021929016311502

8. Collins TD, Ghoussayni SN, Ewins DJ, Kent JA. A six degrees-of-freedom marker set for gait analysis: repeatability and comparison with a modified Helen Hayes set. Gait Posture. 2009 Aug;30(2):173–80.

9. Schmitz A, Buczek FL, Bruening D, Rainbow MJ, Cooney K, Thelen D. Comparison of hierarchical and six degrees-of-freedom marker sets in analyzing gait kinematics. Comput Methods Biomech Biomed Engin. 2015;(August):1–9.

10. Benoit DL, Ramsey DK, Lamontagne M, Xu L, Wretenberg P, Renström P. Effect of skin movement artifact on knee kinematics during gait and cutting motions measured in vivo. Gait Posture. 2006 Oct;24(2):152–64.

11. Fiorentino NM, Atkins PR, Kutschke MJ, Goebel JM, Foreman KB, Anderson AE. Soft tissue artifact causes significant errors in the calculation of joint angles and range of motion at the hip. Gait Posture. 2017 Jun;55:184–90.

12. Stagni R, Fantozzi S, Cappello A, Leardini A. Quantification of soft tissue artefact in motion analysis by combining 3D fluoroscopy and stereophotogrammetry: a study on two subjects. Clin Biomech. 2005 Mar;20(3):320–9.

13. Nisell R, Németh G, Ohlsén H. Joint forces in extension of the knee. Analysis of a mechanical model. Acta Orthop Scand. 1986 Feb;57(1):41–6.

14. Isman RE, Inman VT, Poor P. Anthropometric studies of the human foot and ankle. Bull Prosthet Res. 1969;11:97–108.

15. Lu TW, O’Connor JJ. Bone position estimation from skin marker co-ordinates using global optimisation with joint constraints. J Biomech. 1999;32(2):129–34.

16. Anderson FC, Pandy MG. Static and dynamic optimization solutions for gait are practically equivalent. J Biomech. 2001;34(2):153–161.

17. Schwartz MH, Rozumalski A. A new method for estimating joint parameters from motion data. J Biomech. 2005;38(1):107–16.

18. Rajagopal A, Dembia C, DeMers M, Delp D, Hicks J, Delp S. Full body musculoskeletal model for muscle-driven simulation of human gait. IEEE Trans Biomed Eng. 2016;9294(c):1–1.

19. Delp SL, Anderson FC, Arnold AS, Loan P, Habib A, John CT, et al. OpenSim: Open-Source Software to Create and Analyze Dynamic Simulations of Movement. IEEE Trans Biomed Eng. 2007 Nov;54(11):1940–50.

20. Akbarshahi M, Schache AG, Fernandez JW, Baker R, Banks S, Pandy MG. Non-invasive assessment of soft-tissue artifact and its effect on knee joint kinematics during functional activity. J Biomech. 2010 May 7;43(7):1292–301.

21. Reinschmidt C, Van Den Bogert AJ, Lundberg A, Nigg BM, Murphy N, Stacoff A, et al. Tibiofemoral and tibiocalcaneal motion during walking: External vs. Skeletal markers. Gait Posture. 1997;6(2):98–109.

22. Piazza SJ, Okita N, Cavanagh PR. Accuracy of the functional method of hip joint center location: Effects of limited motion and varied implementation. J Biomech. 2001;34(7):967–73.

23. Daysal GA, Goker B, Gonen E, Demirag MD, Haznedaroglu S, Ozturk MA, et al. The relationship between hip joint space width, center edge angle and acetabular depth. Osteoarthritis Cartilage. 2007 Dec 1;15(12):1446–51.

24. Grood ES, Suntay WJ. A joint coordinate system for the clinical description of three-dimensional motions: application to the knee. J Biomech Eng. 1983;105(2):136–44.

25. Wu G, Siegler S, Allard P, Kirtley C, Leardini A, Rosenbaum D, et al. ISB recommendation on definitions of joint coordinate system of various joints for the reporting of human joint motion--part I: ankle, hip, and spine. International Society of Biomechanics. J Biomech. 2002 Apr;35(4):543–8.

26. Challis JH. A procedure for determining rigid body transformation parameters. J Biomech. 1995 Jun;28(6):733–7.

27. Zeni J, Richards J, Higginson JS. Two simple methods for determining gait events during treadmill and overground walking using kinematic data. Gait Posture. 2008 May;27(4):710–4.

28. Baxter JR, Sturnick DR, Demetracopoulos CA, Ellis SJ, Deland JT. Cadaveric gait simulation reproduces foot and ankle kinematics from population-specific inputs. J Orthop Res. 2016 Jan;1–6.

29. Lenhoff MW, Santner TJ, Otis JC, Peterson MGE, Williams BJ, Backus SI. Bootstrap prediction and confidence bands: A superior statistical method for analysis of gait data. Gait Posture. 1999;9(1):10–7.

30. McGinley JL, Baker R, Wolfe R, Morris ME. The reliability of three-dimensional kinematic gait measurements: A systematic review. Gait Posture. 2009 Apr;29(3):360–9.

31. Murray MP, Mollinger LA, Gardner GM, Sepic SB. Kinematic and EMG patterns during slow, free, and fast walking. J Orthop Res. 1984 Jan 1;2(3):272–80.

32. Zeni Jr. JA, Higginson JS. Differences in gait parameters between healthy subjects and persons with moderate and severe knee osteoarthritis: A result of altered walking speed? Clin Biomech. 2009 May;24(4):372–8.

33. Hurwitz DE, Foucher KC, Sumner DR, Andriacchi TP, Rosenberg AG, Galante JO. Hip motion and moments during gait relate directly to proximal femoral bone mineral density in patients with hip osteoarthritis. J Biomech. 1998;31(10):919–925.

34. Buczek FL, Rainbow MJ, Cooney KM, Walker MR, Sanders JO. Implications of using hierarchical and six degree-of-freedom models for normal gait analyses. Gait Posture. 2010;31(1):57–63.

35. Arampatzis A, De Monte G, Karamanidis K, Morey-Klapsing G. Influence of the muscle tendon unit’s mechanical and morphological properties on running economy. J Exp Biol. 2006;3345–57.

36. Lamberto G. To what extent is joint and muscle mechanics predicted by musculoskeletal models sensitive to soft tissue artefacts? J Biomech. 2017;

37. Li J-D, Lu T-W, Lin C-C, Kuo M-Y, Hsu H-C, Shen W-C. Soft tissue artefacts of skin markers on the lower limb during cycling: Effects of joint angles and pedal resistance. J Biomech. 2017 Sep 6;62:27–38.

38. Elliott G, Sawicki GS, Marecki A, Herr H. The biomechanics and energetics of human running using an elastic knee exoskeleton. In: 2013 IEEE 13th International Conference on Rehabilitation Robotics (ICORR). 2013. p. 1–6.

39. Sawicki GS, Ferris DP. Powered ankle exoskeletons reveal the metabolic cost of plantar flexor mechanical work during walking with longer steps at constant step frequency. J Exp Biol. 2009 Jan;212(Pt 1):21–31.

40. Lichtwark GA, Wilson AM. In vivo mechanical properties of the human Achilles tendon during one-legged hopping. J Exp Biol. 2005 Dec;208(Pt 24):4715–25.

41. Huang C, Chen X, Cao S, Zhang X. Muscle-tendon units localization and activation level analysis based on high-density surface EMG array and NMF algorithm. J Neural Eng. 2016;13(6):066001.

42. Kainz H, Graham D, Edwards J, Walsh HPJ, Maine S, Boyd RN, et al. Reliability of four models for clinical gait analysis. Gait Posture. 2017 Apr 3;54:325–31.

43. Schmitz A, Piovesan D. Development of an Open-Source, Discrete Element Knee Model. IEEE Trans Biomed Eng. 2016 Oct;63(10):2056–67.

44. Kar J, Quesada PM. A Numerical Simulation Approach to Studying Anterior Cruciate Ligament Strains and Internal Forces Among Young Recreational Women Performing Valgus Inducing Stop-Jump Activities. Ann Biomed Eng. 2012;

45. Marques F, Souto A, Flores P. On the constraints violation in forward dynamics of multibody systems. Multibody Syst Dyn. 2017;

46. Nester C, Jones RK, Liu A, Howard D, Lundberg A, Arndt A, et al. Foot kinematics during walking measured using bone and surface mounted markers. J Biomech. 2007;40(15):3412–23.

47. Chehab EF, Andriacchi TP, Farve J. Speed, age, sex, and body mass index provide a rigorous basis for comparing the kinematic and kinetic profiles of the lower extremity during walking. 2017;

48. Kainz H, Modenese L, Lloyd DG, Maine S, Walsh HPJ, Carty CP. Joint kinematic calculation based on clinical direct kinematic versus inverse kinematic gait models. J Biomech. 2016 Jun 14;49(9):1658–69.

49. Kainz H, Carty CP, Maine S, Walsh HPJ, Lloyd DG, Modenese L. Effects of hip joint centre mislocation on gait kinematics of children with cerebral palsy calculated using patientspecific direct and inverse kinematic models. Gait Posture. 2017 Sep;57:154–60.

